# Pangenomic analysis reveals pathogen-specific regions and novel effector candidates in *Fusarium oxysporum* f. sp. *cepae*

**DOI:** 10.1101/182238

**Authors:** Andrew D. Armitage, Andrew Taylor, Maria K. Sobczyk, Laura Baxter, Bethany P.J. Greenfield, Helen J. Bates, Fiona Wilson, Alison C. Jackson, Sascha Ott, Richard J. Harrison, John P. Clarkson

## Abstract

A reference-quality assembly of *Fusarium oxysporum* f. sp. *cepae* (Foc), the causative agent of onion basal rot has been generated along with genomes of additional pathogenic and non-pathogenic isolates. Phylogenetic analysis confirmed a single origin of the Foc pathogenic lineage.

Genome alignments with other *F. oxysporum* ff. spp. and non pathogens revealed high levels of syntenic conservation of core chromosomes but little synteny between lineage specific (LS) chromosomes. Four LS contigs in Foc totaling 3.9 Mb were designated as pathogen-specific (PS). A two-fold increase in segmental duplication events was observed between LS regions of the genome compared to within core regions or from LS regions to the core.

RNA-seq expression studies identified candidate effectors expressed *in planta*, consisting of both known effector homologs and novel candidates. FTF1 and a subset of other transcription factors implicated in regulation of effector expression were found to be expressed *in planta.*

## Introduction

The pan-genome of many microbial species, including bacteria, fungi^1^ and oomycetes^2^ comprises core genes, often in syntenically conserved, gene-dense regions of the genome and so-called ‘dispensable’ genes, located in gene-sparse regions, flanked by abundant transposable elements^3,4^. In *Fusarium* and other fungal phytopathogens these may be specific areas of conserved ‘core’ chromosomes and individual lineage-specific ‘LS’ chromosomes, also known as dispensable or supernumerary chromosomes^1^.

The fungal pathogen *Fusarium oxysporum (Fo)* represents a diverse group of *formae speciales* (ff. spp.) causing crown and root rots as well as vascular wilts^5^. These ff. spp. are part of the *Fo* species complex (containing over 150 members^6^), show narrow host adaptation, and exploit a broad range of niches. Many *Fo* isolates are non-pathogenic soil saprophytes and some have even been exploited as biocontrol agents^7,8^.

Bulb onion (*Allium cepa* L.) is a globally important crop; worldwide production in 2013 was 87Mt^9^. One of the major constraints to production is Fusarium basal rot (FBR), caused predominantly by *F. oxysiporum* f. sp. c*epae* (Foc). Symptoms of infection include seedling damping off, root rot and basal rot which spreads up through the bulb scales ^10^. Like many other *Fusarium* species, Foc produces resilient, long-lived chlamydospores that survive in the soil for many years^10,11^.

In a recent UK study, all highly pathogenic Foc isolates were placed in a single clade (divided into 2 sub-clades) based on sequencing of housekeeping genes including EF1-α, whilst non-pathogenic isolates showed much greater diversity and were placed in multiple clades^12^. This suggests a clonal origin of Foc and is supported by work in Japan where all

*Fo* isolates from bulb onion that were shown to be pathogenic were from the same EF1-α or intergenic spacer region (IGS) clade^13^. Previous studies based on EF1α^14–16^, rRNA^17^, ISSR markers^17^ and RAPD markers^14^ have suggested that Foc is more diverse with isolates falling into multiple clades with a potentially polyphyletic origin. However, such studies are rarely associated with large scale pathogenicity testing and many isolates often prove to be non-pathogenic on onions, even though they have been isolated from diseased plants^12^.

Analysis of tomato plants infected with *Fo* f. sp. *lycopersici* (Fol) has contributed to the identification of 14 *F. oxysporum* proteins that are secreted into the xylem sap (SIX 1-14)^18^ and knockout studies have confirmed that some of these *SIX* genes (e.g. *SIX3*, *SIX5* and *SIX6)* contribute to pathogenicity and therefore code for effector proteins^19,20^. Homologs of *SIX* genes have also been identified in a wide range of *F. oxysporum* ff. spp^21^.

Genome sequencing of Fol isolate 4287 identified LS regions including the ends of chromosomes 1 and 2, as well whole chromosomes 3, 6 14 and 15^1^. These are hypothesised to have been acquired through horizontal gene transfer^1^. Of the four LS chromosomes identified in Fol, chromosome 14 has been characterised as being primarily responsible for conferring pathogenicity. LS regions with a size range of 4-19 Mb have since been identified in a number of other ff. spp. including those infecting cucurbits and legumes and appear to be responsible for host specificity in different ff. spp.^21–23^.

With the exception of SIX13, all putative SIX effectors identified in Fol 4287, are located on Fol chromosome 14, which is also enriched for secreted proteins and secondary metabolite genes^4^. This leads to questions over the function of other Fol LS chromosomes that are not strongly implicated in pathogenicity. Due to the difficulties with assembling effector-containing regions with short-read technologies, it is still unclear whether other ff. spp. possess four LS chromosomes syntenic to those in Fol or unique complements^21^. *Fo* ff. spp. have been shown to possess some effectors that are common to all sequenced *Fo* isolates (including non-pathogens) as well as effector complements specific to each f. sp.^23^. However, the distribution of these ‘core’ effectors throughout the genome has not yet been determined.

Transcriptional regulation of *SIX* genes and *Fo* effector complements is still relatively poorly understood. Nine transcription factors have been identified on Fol LS regions (TF1-9)^24^. Of these, TF1 (*FTF1*), a Zn(II)2Cys6-type transcription factor is the best characterised, and regulates *SIX* gene expression *in planta^24–26^*. However, *FTF1* contains a number of homologous genes within the Fol genome, including within core chromosomes^25^. Transcriptional regulation of LS effector genes is dependent upon factors located on core chromosomes, e.g. *SGE1* is required for differential expression of Fol effector genes *in planta^24,27^*.

*SIX* genes and other putative effectors are closely associated with miniature inverted-repeat transposable elements (MITEs)^4^. Two classes of MITEs are associated with *SIX* genes and are thought to be derived from ancestral transposable elements. Miniature impala (mimp) sequences are found in promoter regions upstream of *SIX* genes 1-14 and mFot5 sequences are located downstream of *SIX* genes 2, 4, 5 and 7. Although they are present in the promoter regions of *SIX* genes, deletion of mimps has not led to differences in *SIX* gene expression *in planta^4^*.

As well as differing between *Fo* ff. spp., *SIX* gene complements can also vary within races of individual f. sp., indicating that there may be loss of genes that do not provide an advantage to pathogenicity.. *SIX*4 has been shown to act as an AVR gene, triggering resistance in tomato through interaction with the *I-1* resistance gene^28^. Fol races 2 and 3 lack *SIX*4 and have restored virulence on *I-1* resistant tomato varieties. It is therefore important to understand the capacity for effector loss in response to selection pressures from deployment of resistant hosts as well as any variation in effector complement in natural populations of pathogenic *Fo*.

The main aim of this study was to investigate functional specialisation of chromosomes within the Foc genome, using pangenomic comparisons of pathogenic and non-pathogenic isolates and *in planta* expression studies.

## Results

### Single molecule sequencing yields a near-complete genome assembly

Three pathogenic Foc (Fus2, 125, A23) and four non-pathogenic *Fo* isolates (A13, A28, PG, CB3) from onion were selected for whole genome sequencing based upon previous work^12^. Highly pathogenic isolates were collected from different UK locations, each containing seven *SIX* genes, whereas the four non-pathogenic isolates had one or no *SIX* genes. The pathogenic Foc isolate Fus2 has been used in multiple experiments assessing isolate virulence and onion resistance^12,15^ and was selected for further sequencing using PacBio long-read technology. (Table 1, section A). The Fus2 assembly yielded a 34 contig, 53.4 Mb reference genome while the remaining isolates yielded *de novo* assemblies of 50-55 Mb in 920-3121 contigs. Gene space within assemblies was shown to be comparable with over 99% of 3725 core Sordariomycete genes (BUSCO) present in all assemblies (Table 1, section A), values comparable to previous *Fusarium* sequencing projects^1^.

**Table 1.** Summarised assembly statistics for *F. oxysporum* (Fo) and F. oxysporum f. sp. cepae (FOC) isolates (A) sequenced as part of this study and summaries of published genomes for the Fo isolate fo47 and F. oxysporum fsp. lycopersici (FOL) isolate 4287. Gene models predicted in this study show values predicted by Braker and additional genes predicted by CodingQuary (B).

| Organism Isolate | FOC Fus2 | FOC 125 | FOC A23 | Fo A13 | Fo A28 | Fo CB3 | Fo PG | Fo fo47<sup>1</sup> | FOL 4287<sup>1</sup> |
| --- | --- | --- | --- | --- | --- | --- | --- | --- | --- |
| **A) Assembly stats:** |  |  |  |  |  |  |  |  |  |
| Total coverage (fold) | 214 | 63 | 47 | 33 | 53 | 30 | 69 | - | - |
| Technology | PacBio + MiSeq | MiSeq | MiSeq | MiSeq | MiSeq | MiSeq | MiSeq | - | - |
| Assembly size (Mb) | 53.4 | 51.4 | 51.0 | 54.8 | 53.1 | 50.5 | 50.3 | 49.7 | 61.5 |
| Contigs | 34 | 2121 | 1999 | 3121 | 2375 | 1720 | 920 | 124 | 15 + 74* |
| Largest contig (Kb) | 6,434 | 573 | 538 | 806 | 1,221 | 1,006 | 2,304 | 6,199 | 6,855 |
| N50 (Kb) | 414 | 128 | 147 | 167 | 297 | 237 | 413 | 3,844 | 4,590 |
| Sodariomycete genes (BUSCO) | 3687 | 3686 | 3688 | 3692 | 3683 | 3679 | 3684 | 3687 | 3599 |
| % Sodariomycete genes (BUSCO) | 99 | 99 | 99 | 99 | 99 | 99 | 99 | 99 | 97 |
| % Repeatmasked | 10.54 | 7.36 | 6.96 | 9.03 | 8.35 | 5.67 | 6.11 | 5.68 | 16.42 |
| **B) Gene models:** |  |  |  |  |  |  |  |  |  |
| Total genes | 18855 | 18505 | 18323 | 18790 | 18629 | 17943 | 17830 | 18191 | 20925 |
| Total proteins | 19371 | 18743 | 18557 | 18934 | 18874 | 18178 | 18101 | 24818 | 27347 |
| Braker transcripts | 17823 | 17197 | 17006 | 17986 | 17426 | 16833 | 16811 | - | - |
| CodingQuary transcripts | 1548 | 1546 | 1551 | 948 | 1448 | 1345 | 1290 | - | - |
| Sodariomycete genes (BUSCO) | 3668 | 3671 | 3663 | 3673 | 3627 | 3663 | 3664 | 3687 | 3577 |
| % Sodariomycete genes (BUSCO) | 98 | 99 | 98 | 99 | 97 | 98 | 98 | 99 | 96 |
| Secreted Genes | 1449 | 1439 | 1431 | 1505 | 1442 | 1433 | 1429 | 1409 | 1493 |

### Gene annotation in the Foc genome

Gene prediction resulted in 17,830 – 18,855 genes in *Fo* and Foc assemblies (Table 1, section B), again comparable to published gene models^1^. Differences in gene prediction approaches were apparent in the number of predicted proteins, with greater numbers of alternative transcripts predicted in fo47 and Fol 4287 genomes (Table 1). BUSCO analysis showed a low false negative rate, comparable to that of fo47 and Fol 4287. Assemblies were submitted as Whole Genome Shotgun projects to DDBJ/ENA/Genbank (Supp. Table 1).

### Phylogeny confirms a single clade of onion-infecting Foc

A *BEAST phylogenetic tree of the thirty single copy genes present in all Eukaryotic fungi for all available *Fo* ff. spp. and non-pathogenic isolates revealed a well-supported clade for all sequenced pathogenic isolates of Foc indicating a monophyletic origin (Figure 1). Non-pathogenic *Fo* strains from onion were interspersed throughout the phylogeny and clustered with a range of other ff. spp. For example, isolate PG from onion is in the clade of Fol 4287.

**Figure 1.**
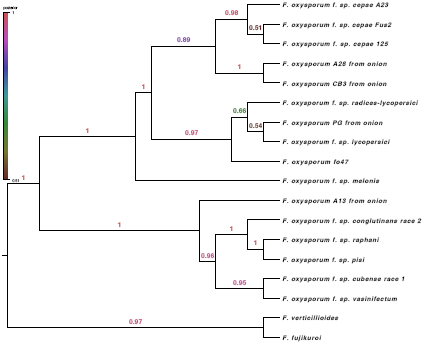
Bayesian phylogeny of *Fusarium oxysporum* isolates from onion and other hosts using 30 single copy loci. Pathogenic Foc isolates (Fus2, 125, A23) are monophyletic within the phylogeny while non-pathogenic isolates from onion (A13, A28, CB3, PG), are interspersed throughout the tree.

### Genome alignment allows identification of pathogen-specific chromosomes and genomic regions

Alignments of the six *de novo* assembled genomes to the Foc Fus2 reference genome allowed identification of core and LS regions between the three pathogenic Foc and three non-pathogenic *Fo* isolates (Table 2). Core contigs represented 46.7 Mb of the Fus2 assembly, while seven LS contigs represented 5.7 Mb of the assembly. Fus2 contig 18 was identified as LS by this analysis; however, later synteny analysis in comparison with Fol showed some conservation of synteny across this contig and it was therefore considered a poorly aligned region of a core chromosome. Of the seven identified LS contigs, four were only found in the three Foc pathogens, representing 3.9 Mb of the assembly, and were designated as “Pathogen Specific” (PS) contigs. The reciprocal alignments of Foc and non-pathogenic *Fo* isolates against the Fol genome clearly identified known Fol LS chromosomes (Supp. Table 2). An additional 11 Foc Fus2 contigs, (440kb in total including mitochondial sequences) were smaller than 200 kb and were excluded from this analysis.

**Table 2.**
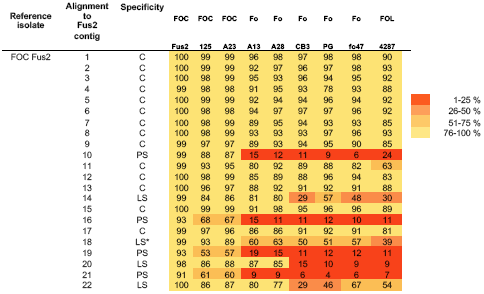
Identification of lineage specific regions in *F. oxysporum* f. sp. *cepae* (FOC) genomes, through alignment of *F. oxysporum* (Fo) and *f. sp. lyccopersici* (FOL) assemblies. The percentage of non>masked bp covered by aligned sequence is shown for each reference contig. * Contig 18 was identified as LS, but later analysis showed synteny to FOL chromosome 12 and alignment of Fo raw sequencing reads across this region.

### Single copy orthologous genes allow mapping of Foc contigs to Fol chromosomes

Foc Fus2 contigs were assigned a chromosome ID consistent with the Fol 4287 assembly following synteny analysis of orthologous genes between Fol 4287 and Foc Fus2. Orthology analysis identified genes common between all *Fo* isolates and those unique to Foc and Fol; 97% proteins were clustered into 15607 orthogroups, 10891 orthogroups were shared between all *Fo* isolates and represented 151,631 proteins while 7396 orthogroups contained proteins in a 1:1 relationship between Foc Fus2 and Fol 4287. The location of genes encoding these proteins allowed macrosynteny to be assessed between the 15 Fol 4287 chromosomes and the 34 assembled Foc contigs. Of the 34 Foc contigs, 12 were smaller than 200 kb and were considered too small to be assigned. Of the remaining 22 Foc contigs, 15 were assigned to the 11 core Fol chromosomes (Figure 2). The remaining 7 Foc contigs did not show clear synteny with Fol chromosomes, supporting their designation as LS in the alignment analysis presented above.

**Figure 2.**
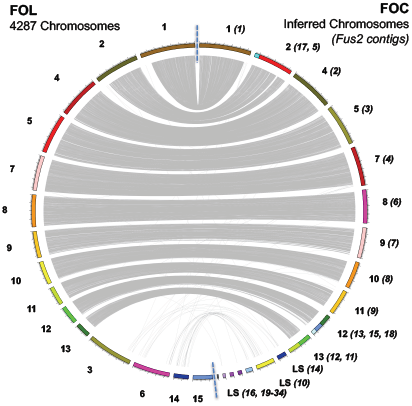
Synteny of chromosomes between Fol and Foc genome assemblies. Relationships are shown through linking single copy orthologous genes, present in both genomes. Core chromosomes can be identified through synteny between Foc and Fol whereas LS regions show reduced synteny. The number of LS chromosomes does not appear to be conserved between assemblies. Fol chromosome 15 harbours no genes in single copy orthogroups.

### Effector annotation of the Fus2 genome

Generic effector-prediction approaches based upon identification of secreted proteins with an effector-like structure (small, cysteine rich), secreted carbohydrate active enzymes (CAZymes) (Table 3, section A) and identification of secondary metabolite synthesis genes (Table 3, section B) were used to investigate Foc and *Fo* effector complements. Secreted proteins represented 7.6 % of the Fus2 proteome, and genes encoding proteins with an effector-like structure approximately 1.9 % of the proteome. Overall, comparisons between genomes showed no significant differences between the total numbers of Foc and non-pathogenic *Fo* genes encoding secreted proteins (t = 0.23, df = 4.8, p-value = 0.83), EffectorP proteins t = −1.29, df = 4.02, p-value = 0.26, or secreted CAZymes (t = −0.72, df = 4.07, p-value = 0.51).

**Table 3.** Summarised assembly statistics for *F. oxysporum* (Fo) and *F. oxysporum f. sp. cepae* (FOC) isolates with *F. oxysporum f. sp. lycopersici* (FOL) for comparison. Number of genes predicted as secreted proteins (A), or within 2 Kb of a mimp sequence are reported (B) along with numbers of secondary metabolite clusters (C).

| Organism<br>Isolate | FOC<br>Fus2 | FOC<br>125 | FOC<br>A23 | Fo<br>A13 | Fo<br>A28 | Fo<br>CB3 | Fo<br>PG | Fo<br>fo47<sup>1</sup> | FOL<br>4287<sup>1</sup> |
| --- | --- | --- | --- | --- | --- | --- | --- | --- | --- |
| **A) Effector candidates:** |  |  |  |  |  |  |  |  |  |
| Secreted & effector-like structure | 355 | 357 | 355 | 364 | 346 | 357 | 337 | 291 | 351 |
| Secreted CAZYmes | 386 | 386 | 387 | 397 | 376 | 383 | 381 | 382 | 386 |
| **B) Secondary metabolites:** |  |  |  |  |  |  |  |  |  |
| Gene clusters | 50 | 46 | 46 | 49 | 48 | 44 | 45 | 47 | 49 |
| Genes in clusters | 703 | 559 | 561 | 626 | 656 | 604 | 638 | 705 | 701 |
| **C) Mimps** |  |  |  |  |  |  |  |  |  |
| mimps in genome | 153 | 140 | 136 | 55 | 35 | 30 | 51 | 25 | 142 + 16* |
| Genes in 2kb of Mimp | 155 | 95 | 88 | 49 | 36 | 32 | 43 | 24 | 108 |
| Secreted genes in 2kb of Mimp | 31 | 24 | 20 | 9 | 5 | 1 | 3 | 3 | 22 |

### Mimp distribution in the Fus2 genome and prediction of mimp-associated effector candidates

Mimp sequences are often found in proximity to *Fo* effectors^4^. Foc genomes were found to contain 136-153 mimps, significantly more than the 25-55 observed in non-pathogenic *Fo* isolates (t = −13.29, df = 5.75, p-value < 0.01) (Table 3, section C). Of the 153 mimps within the Foc Fus2 genome, the majority were distributed throughout LS regions, with 120 in newly designated PS regions and 20 in non-PS LS regions. Of the 158 mimps identified in the Fol genome^4^, 132 of these were present in Fol LS chromosomes, 10 in Fol core chromosomes and 16 in unplaced Fol contigs.

### Foc core chromosomes 11-13 are enriched for secreted proteins and cell wall degrading enzymes

Foc core chromosomes 11-13 were noted to contain many genes with an effector-like structure and secreted CAZymes (Figure 3). Comparison of the density of genes between core, effector-rich core (chromosomes 11-13), non-PS LS and PS regions of the genome (Figure 4) showed that secreted genes (SignalP) were present at different densities between core, effector-rich core, LS and PS regions of the Fus2 genome (F(3,14) = 8.072, P < 0.01). Similarly, the density of secreted CAZymes (F(3,14) = 12.04, P < 0.01), EffectorP genes (F(3,14) = 7.506, P < 0.01) and secondary metabolite clusters (F(2,12) = 4.26, P = 0.04) also differed between these regions. Effector-rich core regions were found to contain secreted genes at 50 genes Mb^−1^; double the density at which these genes were found in other regions of the genome (19-24 genes Mb^−1^). Similarly, secreted CAZYmes were highly enriched within this region at 16 genes Mb^−1^ in comparison to 3-6 genes Mb^−1^ in other regions. The effector-rich core also showed high densities of EffectorP genes and secondary metabolite clusters, although these were not significantly enriched in comparison to core regions (Figure 4).

**Figure 3.**
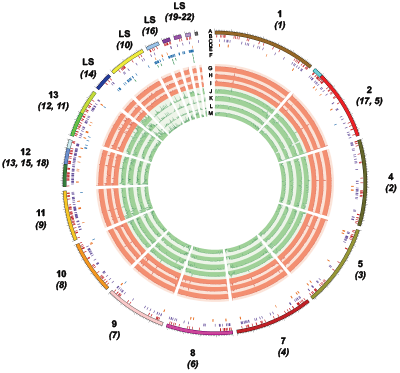
Visualisation of Foc isolate Fus2 genome assembled into 34 contigs (A). Locations of predicted secreted effectors (B), secreted carbohydrate active enzymes (C), secondary metabolite gene clusters (D), mimp sequences (E) and SIX gene homologs (F) are identified within contigs. Alignment of assemblies from pathogenic Foc isolates Fus2, 125 and A23 (G-I), non-pathogenic isolates A28, PG, CB3 and A13 (J-M) and *F. proliferatum* isolate A8 (N) are shown.

### Identification of effector candidates in PS regions

Fus2 PS regions contained 34 secreted EffectorP genes, 11 secreted CAZymes and 4 secondary metabolite gene clusters (Supp. Table 3), of which 14, 4 and 3 were within 2 kb of a mimp respectively. Non-PS LS regions contained an additional 6 EffectorP genes and 7 Secreted CAZymes; in each case two of these were within 2 kb of a mimp. Two of the four PS secondary metabolite gene clusters were encoded a terpene synthesis gene cluster and one encoded a polyketide synthesis gene cluster, none of which were within 2 kb of a mimp.

The Foc genomes were confirmed to carry homologs of the seven Fol *SIX* genes (SIX3, 5, 7, 9, 10, 12 and 14) through BLAST searches (Supp. Table 4) as reported previously^12^ with two homologs of *SIX3* and *SIX9* present. All *SIX* genes identified within the Foc Fus2 genome were located within the four PS contigs 10, 16, 19 and 21 (Supp. Table 5, Figure 3). Searches for *SIX* genes in the non-pathogenic *Fo* isolates from onion also confirmed that isolate PG contained *SIX9^12^*.

Investigation into sequence conservation between Foc isolates found no non-synonymous variation between EffectorP genes and secreted CAZymes in PS regions. Similarly, very little variation was seen in these effector candidates in other regions of the genome, with only five EffectorP genes and secreted CAZymes showing variation between Foc isolates (Supp. Table 6).

### Effector candidates in LS regions show lower codon usage bias than those located in the core genome

Codon usage was investigated across Foc Fus2 genes in core and LS regions (Supp. Table 7). Mean CBI values for secreted CAZY and secreted EffectorP candidates were 0.385 and 0.336 respectively, significantly higher (*t*-test, p-value < 10^−18^) than found for all genes (mean CBI=0.215). In the Fol genome, Ma *et al.* (2010) showed a shift in preferred codons between LS and core chromosomes, especially towards GC in their third position^1^. In Foc there was no change in GC content across these two portions of the genome (core chromosomes, mean 51.7%; LS chromosomes 51.3%). LS regions (mean CBI=0.145) contained genes with a significantly lower (*t*-test, p-value< 2.2e-16) average codon usage bias than in the core genome (mean CBI=0.221). All genes with close proximity (< 2 kb) to mimps showed relaxed codon usage bias and LS secreted genes showed significant (t-test, p-value=8.84E-03) codon usage bias (mean CBI=0.284) compared to the rest of the LS genes (mean CBI=0.145).

### Foc LS regions have higher levels of gene duplication than core chromosomes

Many orthogroups on Fol and Foc LS regions contained inparalogs, with homologous genes elsewhere in LS regions (Supp. Figure 1). Duplicated genes in LS regions showed a bias towards being shared with other LS regions, whereas core Foc regions exhibited lower levels of gene duplication, with a concentration of duplicated genes shared with terminal regions of other chromosomes/contigs. In total, 1,424 gene clusters, representing one or more duplication events were identified. Taking one representative focal gene in each cluster and comparing the chromosomal/contig location between pairs of genes in these clusters, 49 % of duplication events were observed to be within or between LS chromosomes/contigs (3,186). In contrast, 28 % of duplications were between core and LS (1,822) and 23 % within or between core and core (1,517) chromosomes. Overall, 1,235 of 2,115 genes on LS contigs contained a paralog in the genome, corresponding to a lower density of duplicated genes on core than on LS chromosomes (permutation test, *p*-value <0.001). Dissection of duplications into tandem and segmental did not reveal any further patterns, with only 69 tandem duplications identified.

### Open reading frame density is maintained in Foc LS regions

As in Fol, Foc LS contigs showed increased levels of repetitive and low complexity content in comparison to core regions, with 3-15% repeat-masked in core regions compared with 33-59 % in LS regions respectively (Supp. Figure 2). Gene density of Fol LS regions has previously been shown to be lower than on core regions^1^ but this was not the case for Foc, where gene density was maintained between core and LS regions (Supp. Fig. 2). Analysis of predicted gene function for Foc LS regions found that Interproscan terms associated with transposon activity were significantly enriched on non-PS LS regions and PS regions (Supp. Table 8, P < 0.05). Terms associated with Helitron helicase transposons were found exclusively on PS regions. Aside from genes with transposon-associated features, PS regions showed enrichment for genes lacking IPR annotations (Supp. Table 8), a feature of newly described gene families.

### Known effectors and effector candidates in PS regions are among the highest expressed genes *in planta*

Using an established *in vitro* onion seedling root infection system, expression of Foc genes *in planta* was explored at 72 hours post inoculation (hpi), a previously identified timepoint when *SIX* genes are highly expressed^12^. Proximity to a mimp was associated with greater expression, with both non-effector genes and secreted genes within 2 kb of a mimp showing significantly greater expression than similar non-mimp genes (Figure 5). Genes with a putative effector status (using the EffectorP pipeline) in PS regions showed high expression *in planta*, irrespective of proximity to a mimp.

Foc genes showing the highest expression *in planta* were further investigated. 21 LS Fus2 genes had equal or greater expression than the 50 highest expressed core genes (Supp. Table 9), with 17 of these on PS regions. These 17 highly expressed PS genes included 11 genes predicted as both secreted and within 2 kb of a mimp and represented previously-identified and novel effector candidates.

Foc PS effector candidates, *SIX*5 and two *SIX*3 homologs were the three highest expressed genes. Additional *SIX* gene homologs also showed high expression *in planta*, with two *SIX*9 homologs the 7^th^ and 17^th^ highest expressed genes. Other *SIX* homologs showed lower levels of expression *in planta*, with *SIX*10, *SIX*7 and *SIX*14 ranking as the 159^th^, 137^th^ and 3277^th^ highest expressed genes, respectively.

Other PS effector candidates represent putative novel effectors. They did not possess any recognisable interproscan domains and did not have an orthologous gene in the Fol gene models. tBLASTx searches against the PHIbase database did not identify any homologs for these genes (e < 1 x 10^−10^) and many showed no homology to sequences on NCBI (e < 1 x 10^−10^). Apart from the 11 genes predicted as both secreted and within 2 kb of a mimp, six additional genes were present on PS regions and also highly expressed. Five of these, including *SIX*12, were located within 2 kb of a mimp and one carried a peptidase domain (IPR001506) indicating that some of these genes may represent additional effector candidates. However, two genes were annotated as membrane-bound proteins discounting them as effector candidates. tBLASTx searches against the PHIbase database did not identify any homologs for these genes (e < 1x10^−10^) and searches against genbank (e < 1 x 10^−10^) found a homolog to only one gene, a hypothetical protein in *Fusarium verticillioides* (BLASTn; e = 1x10^−58^) with 83% identity to the query sequence.

### Non-PS, LS regions of the genome include genes highly expressed *in planta*

In general, genes on non-PS, LS regions showed similar patterns of expression to genes in core regions of the genome (Figure 5). However, four genes on non-PS, LS regions had greater or equal expression to the top 50 expressed genes from core regions of the genome (5^th^, 8^th^, 10^th^ and 11^th^ highest expressed genes, Supp. Table 9). These genes were not considered to be effectors (based on mimps, secretion signals, EffectorP or CAZY identification) but were noted to encode three proteins carrying domains associated with formaldehyde activating enzymes (IPR011057, IPR006913) and a polyketide synthase protein (IPR020843) and were found located in close proximity to one another (g15699-g15700, g15704). As such, they may represent a secondary metabolite cluster not identified by Antismash prediction. Further investigation found that genes carrying a formaldehyde-activating domain (IPR006913) were significantly enriched in non-PS LS contigs with eight proteins identified, in contrast to one on PS regions and 15 on core regions (Supp. Table 8b).

### Helitrons may play a role in re-arrangement of PS regions

Non-canonical Helitrons appear to be a key feature of *Fo* pathogenicity chromosomes^29^. In our analysis only Foc PS regions, were enriched for Helitron helicase-like domains (IPR025476) (Supp. Table 8, panel A) while non-PS LS regions of the genome were not (Supp. Table 8, panel B), with no annotations found in genes on core regions. A total of 35 genes were identified as helitron/helicase-like transposable elements (IPR025476, PF14214) in the Foc genome, with one on an unplaced contig (contig 33) and the rest located on PS regions. Furthermore, these transposable elements showed evidence of expression *in planta*, with six of the 35 genes having a mean fpkm > 5. These results support recent identification of helitrons in *F. oxysporum* ff. spp^29^, and indicate that a Helitron-based mechanism of genomic rearrangement is characteristic of PS regions, rather than core regions, or non-pathogen associated LS regions of the Foc genome.

### Transcription factor analysis in LS regions

To investigate the diversity of transcription factors on Foc LS regions, Interproscan functional annotations were searched for transcription factor domains. This led to the identification of 34 putative transcription factors (Supp. Table 10). Of these, 17 transcription factors were located on non-PS LS regions, of which 12 showed evidence of expression *in planta,* including a homolog of TF4 (Supp. Table 11b). The remaining 17/34 putative TF’s were located on PS regions, and included homologs to previously identified Fol transcription factors TF1, TF3, TF8 and TF9 (Supp. Table 11a). Five PS TF’s showed evidence of expression *in planta* at 72 hpi including a homolog to TF1 (FTF1). With the exception of TF3, each of the previously described TF1-9 genes had a homolog in the core genome, that showed evidence of expression *in planta.*

### Foc carries a distinct complement of FTF1 genes

Due to the association of TF1 (*FTF1*) with *SIX* gene expression in several *Fo* ff. spp.^24^, the FTF gene family was further investigated for all the sequenced *Fo* isolates from onion. A single orthogroup was found to contain all FTF genes previously described in *Fo* strains, fo47 and Fol-4287^25^ as well as the BLAST homologs of FTF genes in Foc and Fo. Alignment and phylogenetic analysis of these genes allowed separation of FTF genes in *FTF1* and *FTF2* families (Figure 6). All Foc isolates were found to carry a single copy of *FTF2*, which was located in the core genome of Fus2. Two copies of *FTF1* were found in Fus2, which were both present in PS contig 19. The two other Foc isolates (125, A23) carried an identical *FTF1* gene in a single copy. Interestingly, *FTF1* homologs were identified in non-pathogenic *Fo* isolates A13, PG and fo47, although these gene sequences were distinct from those in Foc (Figure 6).

## Discussion

In this study, through the use of single molecule sequencing, the structural conservation of lineage-specific (LS) chromosomes has been revealed in the onion basal rot pathogen *F. oxysporum* f. sp. *cepae* through comparison with *F. oxysporum* f. sp. *lycopersci*. Using a pan-genomic approach, this work has revealed for the first time that LS chromosomes can be subdivided into pathogen-specific (PS) and nonspecific chromosomes. RNAseq has shown that known effector candidates, conserved in other *Fusarium* ff. spp., as well as novel effector candidates are expressed *in planta,* paving the way for future functional studies and effector-informed resistance breeding approaches.

Previous studies have indicated that many other ff. spp., including Fol, have a polyphyletic origin^30^ with some exhibiting clear patterns of ‘race’ evolution through stepwise loss of effectors^31^. All pathogenic Foc isolates formed a single clade, consistent with a single origin of pathogenicity on onion^12^. Examination of other non-pathogenic *Fo* isolates from onion showed that these were interspersed throughout the phylogenetic tree, with two isolates forming a sister clade to Foc but other strains, grouping with diverse pathogenic ff. spp. Foc may therefore represent one of the minority of *Fo* ff. spp. With a monophyletic origin, alongside *Fo* f. sp. *ciceris*, f. sp. *canariensis* and f. sp. *albedinis*..

Despite the clear conservation of synteny between Foc and Fol core chromosomes, synteny was not conserved between Foc and Fol LS regions. Interestingly, the LS regions identified in Foc appeared to be much smaller than in Fol, with a total of 5.7 Mb identified in Foc versus 14 Mb in Fol. Designation of 3.9 Mb of Foc LS contigs as PS regions supports previous findings that Foc *SIX3* is located on a ∼ 4 Mb chromosome^13^, indicating that a single pathogenicity chromosome is present in Foc.

Subdivision of LS regions of the Foc genome into PS and non-PS regions showed for the first time that non-pathogenic isolates contain certain LS elements and their presence is independent of the position of the isolate in the phylogenetic tree. For example, contig 14 present in all pathogenic Foc strains, appears to be conserved in non-pathogenic *Fo* isolates A13 and A28 (distantly separated in the phylogeny), but not in isolate CB3, which is most closely related to A28. This suggests that there may be genetic exchange between divergent lineages of non-pathogens in their LS chromosome complement. The functional significance of these non-pathogen associated LS chromosomes is a topic for further investigation.

Structural analysis revealed chromosome-specific patterns of gene duplication, with most gene duplications being segmental and within LS regions. This was consistent with the breakdown of synteny observed between Foc and Fol LS regions. Transposon activity has previously being shown to drive evolution of LS regions in other fungal pathogens^32^. The finding that Helitron-containing transposons are restricted to PS chromosomes suggests that they may be important for the high degree of rearrangements within PS chromosomes. This may be evolutionarily advantageous in a clonal organism as it may facilitate rapid more adaptation, not only through a higher rate of large-scale genomic deletions (which may be adaptive when evading host recognition) but also in preventing the accumulation of linked deleterious mutations from interfering with selection, the so-called Hill-Robertson effect^33^. One question which requires further study, is how helitron helicase transposons are limited to PS chromosomes given there is some evidence for expression.

Similar numbers of effector candidates were identified in both pathogenic and non-pathogenic *Fo* isolates from onion, mainly due to the large number of secreted proteins present in the core genome and on accessory chromosomes, irrespective of pathogenicity. Despite being conserved in *Fo,* Fol chromosome 12 has been reported as conditionally dispensable^34^. A concentration of effector-like genes on the homologous Foc chromosome indicates a functional distinction between Foc effector-rich core chromosomes and the remaining core genome. Foc contains a large complement of core effectors, with enrichment on core chromosomes 11, 12 and 13 (Figure 4). However, these core effector genes were not highly expressed *in planta*. LS contigs present in non-pathogenic isolates from onion contained far fewer mimp sequences, at a level similar to the core genome, far below the levels seen in PS contigs, again highlighting the specific differences between PS chromosomes and the rest of the LS and core genomes.

**Figure 4.**
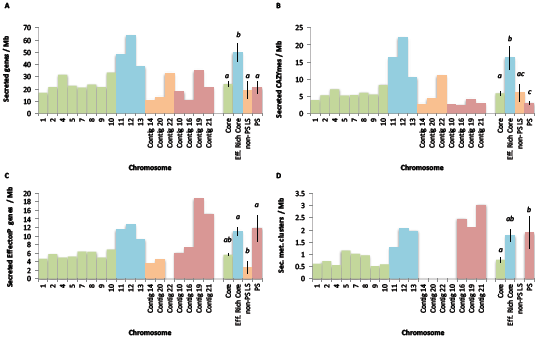
Density of genes associated with an effector-associated function in Foc genomic regions including those encoding: secreted proteins (A); secreted carbohydrate active enzymes (CAZYmes) (B); proteins with an effector-like structure (EffectorP) (C); secondary metabolite gene clusters (D). Average gene density (± SE) is also shown by genomic region including significant differences in gene density by region (ANOVA, P < 0.05).

**Figure 5.**
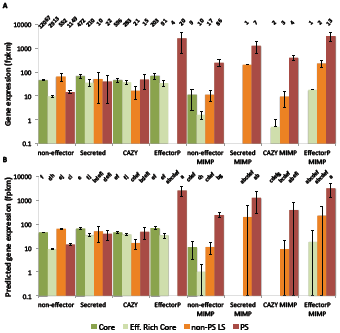
Observed (A) and predicted (B) expression values (mean fpkm) for Foc Fus2 genes expressed during infection of onion seedlings at 72 hpi. Differences in gene expression are observed between effector-type, genomic region and presence of a mimp within 2 kb of the gene. Number of genes in each category is shown above observed values. Pairwise significances (P < 0.05) are shown above predicted values, as determined by a Tukey test of terms from a negative binomial GLM. Effector categories include genes encoding non-effectors, secreted proteins, secreted carbohydrate active enzymes (CAZY) and secreted proteins with an effector-like structure (EffectorP). Genomic regions shown are core chromosomes 1-10 (Core), effector-enriched core chromosomes 11-13 (effector-enriched core), non-PS LS contigs, and PS contigs. Expression is shown for genes within 2 kb of a mimp sequence (mimp).

**Figure 6.**
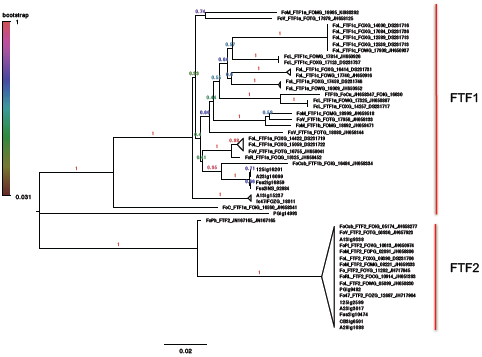
Neighbour joining phylogeny of FTF gene sequences from Foc, Fol, f. sp. *pisi* (FoPi), *radicis-lycopersici* (FoRL), *cubense* (Focub), *vasinifectum* (FoV), *melonis* (FoM), *conglutinans* (Foco) and *phaseoli* (FoPh). Foc FTF1 homologs are distinct from those from other *Fo* ff. spp and in non pathogenic isolates PG and A13. Branches are labelled by bootstrap support from 1000 replicates.

Effector genes on Foc PS regions showed high levels of expression *in planta*; the highest expressed genes were homologs to known *SIX* (most notably *SIX3* and *SIX5*) genes but also included novel effector candidates. It will be important to test whether the I-2 resistance gene, identified in tomato recognises the Foc variants of *SIX3^35^*. Additional non-PS LS regions were also identified; these also possessed high numbers of mimps, and included highly expressed genes.

Transcription factors have previously been identified in the PS chromosome 14 of Fol (13, representing nine gene families;TF1-9)^24^. Foc was found to carry homologs to each of these previously identified TF1-9 genes distributed throughout the core and LS regions, but with no homolog of TF3 in the core genome^24^. However, similar to the different complements of *SIX* genes between *Fo* ff. spp., the pattern of TF1-9 genes present on PS regions in Foc was distinct from Fol. Identification of homologs to known TFs regulating pathogenicity indicates a conservation of transcriptional regulation between Fol and Foc. The role of novel TF candidates on PS regions requires further investigation, including their conservation through comparisons with other *Fo* ff. spp..

One of the major objectives of characterising the genomic basis of pathogenicity is to inform resistance breeding approaches using information about the effector complement of the pathogen^36^. The durability of a single resistance gene is dependent upon the necessity of the detected effector for the infection process and the adaptive potential of the effector gene. Effectors can either mutate to evade recognition by an R gene, such as *SIX3* (*AVR2*) in Fol race3, or be lost such as *SIX4* in Fol races 2 and 3^28^. It is only through the characterisation of effector function, combined with a population genetics approach, that an assessment can be made about the long-term utility of any R gene based resistance in breeding, or the breeding approach needed to effectively deploy the available resistance.

It is therefore important that future work addresses the functional essentiality of effectors, the global diversity in the Foc population, the ability for effectors to mutate and evade recognition and the extent of R gene based resistance in onion.

## Methods

### DNA extraction, library preparation and sequencing

DNA was extracted from freeze-dried mycelium for the three FoC (Fus2, 125, A23) and four *Fo* isolates (A13, A28, PG, CB3) using the Macherey-Nagel Nucleospin Plant II kit (Fisher 11912262). DNA was sheared using the Covaris M220 with microTUBE-50 (Covaris 520166) and size selected using the Blue Pippin (Sage Science). Illumina libraries were constructed using either Illumina TruSeq LT kit (FC-121-2001), or with a PCR-free method using NEBNext End Repair (E6050S), NEBNext dA-tailing (E6053S) and Blunt T/A ligase (M0367S) New England Biolabs modules. Libraries were sequenced using Illumina Miseq v2 2 x 250 bp PE (MS-102-2003) or v3 2 x 300 bp PE (MS-102-3003). Pacbio libraries were prepared by the Earlham Institute UK according to manufacturer specifications and sequenced to achieve approximately 65 times coverage using P6-C4 chemistry. The sequencing of our standard highly pathogenic Foc isolate Fus2 resulted in 69 times and 145 times coverage from PacBio and MiSeq reads, respectively; 30-69 times coverage was generated for the six remaining *Fo* isolates using MiSeq sequencing.

### Genome Assembly

PacBio reads for Foc isolate Fus2 were assembled using Canu and polished using Illumina MiSeq reads in Pilon to correct erroneous SNPs and InDels^37,38^. *De novo* assembly of MiSeq data for the remaining six genomes was performed using Spades v.3.5.0^39^. In all cases Quast^40^ was used to summarise assembly statistics and BUSCO^41^ used to assess completeness of gene space within the assembly. Assemblies were edited in accordance with results from the NCBI contamination screen (run as part of submission to Genbank in November 2016) with contigs split, trimmed or excluded as required. RepeatModeler, RepeatMasker and transposonPSI were used to identify repetitive and low complexity regions (http://www.repeatmasker.org, http://transposonpsi.sourceforge.net). In addition to generating *de novo* assemblies, Illumina sequencing reads were mapped to the PacBio Fus2 assembly. Alignment was performed using Bowtie2 v.2.2.4 before bedtools-intersect was used to determine number of reads aligning across 100 kb windows in Fus2 contigs^42,43^.

### Whole-genome phylogenetic analysis

A thirty gene phylogenetic tree was constructed using selected single copy genes present in the BUSCO ver. 1.22 Eukaryota fungi list for all the *Fo* isolates sequenced in this study as well as additional publically available *Fusarium* spp. genomes^41^. Additional genomes were: (non-pathogenic) *F. oxysporum* (FO_Fo47_V1), *Fo* f. sp. *lycopersici* (FO_MN25_V1), *Fo* f. sp. *conglutinans* (FO_PHW808_V1), *Fo* f. sp. *cubense* (Foc1_1.0), *Fo* f. sp. *melonis* (FO_melonis_V1), *Fo* f. sp. *pisi* (FO_HDV247_V1), *Fo* f. sp. *radices-lycopersici* (FO_CL57_V1), *Fo* f. sp. *raphani* (FO_PHW815_V1), *Fo* f. sp. *vasinfectum* (FO_Cotton_V1), *F. fujikuroi* (assembly EF1), *F. verticillioides* (ASM14955v1) available from EnsemblGenomes Fungi database^44^. CDS sequences of single copy genes conserved across Fungi identified by BUSCO ver. 1.22^41^ that were found to be complete and single copy in all the genome assemblies from this study were used as input in preparing the BEAST phylogenies. In total, 652 such genes were identified and aligned with MAFFT ver. 7.222^45^. The alignments were inspected visually with MEGA7^46^, trimmed and 30 genes in the top 5% highest nucleotide diversity selected for further phylogenetic analysis. A best-fit sequence evolution model for each gene was determined with PartitionFinder ver. 1.1.1^47^ using BIC (Bayesian Information Criterion). The gene trees derived from the set of 30 single copy genes were investigated with multi-locus analysis in *BEAST ver. 2.4.2^48^. For each replicate *BEAST run, 300 million MCMC iterations, sampled every 10,000 chains, were run due to the size of the dataset. The molecular clock was set to strict due to intra-specific sampling and consequent expectation of lower inter-branch rate variation^49^. A Yule prior was placed on species tree and population size model set to follow linear growth with a constant root. The first 10% of results was discarded as burn-in, and run convergence and stationarity were inspected in Tracer ver 1.6 (http://tree.bio.ed.ac.uk/software/tracer/) to confirm that ESS scores for all estimated parameters reached at least stable 200. This was followed by generation of maximum clade credibility tree with median heights with BEAST’s TreeAnnotator and visualisation of the species trees in FigTree ver. 1.4 (http://tree.bio.ed.ac.uk/software/figtree/). Run convergence was established by performing three independent runs and checking the variability of results across runs; a single representative run is reported here.

### Identification of LS regions

LS regions were identified in the Foc Fus2 genome and the previously characterised Fol 4287 assembly using MUMmer v3.23 (PROmer –mum, delta-filter -g)^50^. Repeat-masked assemblies of non-pathogenic Fo, Foc and Fol isolates were aligned against the repeat-masked reference genome^51^. The percentage of unmasked bp covered by aligned sequence was calculated for each reference contig and a threshold of 30% identity was set as the boundary at which a contig was described as present or absent in a strain.

### *In planta* RNAseq

RNAseq data was used to aid gene prediction and assess expression of effector candidates. Onion seedlings were inoculated with either the standard pathogenic Foc Fus2 or the non-pathogenic Fo47 isolate using a sterile, square petri dish system as previously reported^12^. Three replicate plates were set up for each isolate and RNA was extracted from pooled root samples taken from five plants at 72 hpi. Library preparation was carried out using a TruSeq RNA Sample Prep kit V2 (Illumina) and RNA sequencing carried out using an Illumina HiSeq machine with 100 bp paired end reads. Samples were multiplexed over two runs to give approximately 50 million reads per sample.

### Gene prediction

RNAseq reads were aligned to *de novo* assembled genomes using Bowtie2 v.2.2.4 and Tophat v.2.1.0 to aid training of gene prediction programs^42,52^. An initial RNAseq alignment was used to estimate “mean insert size” and “fragment length distribution” of RNAseq reads and Tophat alignments re-run using these parameters. Gene prediction was performed on softmasked genomes using Braker1 v.2^53^, a pipeline for automated training and gene prediction of AUGUSTUS 3.1^54^. Additional gene models were called in intergenic regions using CodingQuary v.2^55^. Braker1 was run using the “fungal” flag and CodingQuary was run using “pathogen” flag.

### Orthology Analysis

Orthology was identified between predicted proteins from the Foc and *Fo* isolates sequenced in this study, the publicly available genomes for *F. oxysporum* isolate Fo47 and Fol isolate 4287^1^. OrthoMCL v.2.0.9^56^ was run with an inflation value of 5 on the combined set of 193,973 predicted proteins from Foc (Fus2, 125, A23), Fo (fo47, A13, A28, PG, CB3) and Fol (4287) genomes. Venn diagrams visualising genes common between Foc and Fo were plotted using the R package VennDiagram^57^.

### Distribution of Duplicated Genes

CDS sequences (one representative longest transcript) of the Foc Fus2 genome were searched against each other using BLASTN, using an E-value threshold of e-10. In order to focus on only recent, lineage-specific duplication events, the BLAST results were filtered to retain only hits with 90% minimum percent identity and minimum 80% subject coverage, which are more stringent criteria than previously applied in similar analyses^58,59^. The BLAST output was subsequently parsed using DAGChainer^60^ in order to identify the reciprocal BLAST hits. Dependent on the position of the genes on the chromosomes, duplications were classified as either tandem (proximal) or segmental (distal). Two parameters: max. distance and number of intervening genes were tested to help classify the duplications as either tandem or segmental and the density of duplication events on chromosomes was plotted using karyoploteR ver. 0.99.8 R package (http://bioconductor.org/packages/devel/bioc/html/karyoploteR.html). Enrichment of duplications in different regions of the genome was tested using permutation tests with 1000 iterations using the regioneR ver. 1.6 R package^61^. In the initial analysis, more than half of identified duplications contained genes with transposon-related InterProScan domains (IPR000477, IPR012337, IPR018289, IPR006600, IPR000477, IPR025476, IPR008906, as well as keywords “transpos*” and “integrase”) and these were subsequently removed from the analysis.

### Functional Annotation and Effector Prediction

Draft functional annotations were determined for gene models using InterProScan-5.18-57.0^62^ and through identifying homology between predicted proteins and those contained in the July 2016 release of the SwissProt database^63^ (using BLASTP (E-value > 1x10^−100^). Interproscan terms were used to test for enrichment of functional domains within PS and non-PS LS regions. Abundance of each interproscan term was tested using Fisher’s exact test, comparing number of genes carrying the annotation to those without. Benjamini Hochberg correction was applied for multiple testing.

Putative secreted proteins were identified through prediction of signal peptides using SignalP v.4.1 and removing those predicted to contain transmembrane domains using TMHMM v.2.0^64,65^. Additional programs were used to provide additional sources of evidence for effectors and pathogenicity factors. EffectorP v1.0 was used to screen secreted proteins for characteristics of length, net charge and amino acid content typical of fungal effectors^66^. Secreted proteins were also screened for carbohydrate active enzymes using HMM models from the CAZY database^67^ and HMMER3^68^. Regions of the genome containing secondary metabolite gene clusters were identified using the Antismash 3.0 webserver^69^. Locations of gene clusters were parsed to gff3 format before genes intersecting these regions were identified using Bedtools.

Genes within 2 kb of a mimp sequence were identified using the consensus sequence for the mimp 3’ inverted repeat^4^. This was searched against assembled genomes using Perl regular expressions /CAGTGGG..GCAA[TA]AA/ and /TT[TA]TTGC..CCCACTG/. Genes within 2 kb of these mimp sequences were marked as candidates for being under the influence of a mimp-containing promoter.

Differences in total numbers of genes encoding secreted proteins, secreted CAZYmes and secreted EffectorP proteins between FoC and Fo isolates were assessed using t-tests in R. Differences in density of genes encoding secreted proteins in different regions of the Fus2 genome were tested using ANOVA in R, with pairwise differences between regions assessed using t-tests with bonferroni correction. Identical analysis were performed to assess density of secreted CAZYmes, secreted EffectorP proteins and density of secondary metabolite clusters by genomic region.

### Codon Bias Amongst Putative Pathogenicity-related Genes

Multivariate correspondence analysis of codon usage to detect presence of codon bias was first investigated using codonW ver. 1.3 (http://codonw.sourceforge.net/). Prior to codonW runs, Foc Fus2 coding sequences (one longest representative sequence per gene) were filtered to remove genes with potentially unusual codon usage stemming from their foreign origin which could indicate mis-annotated false positives; transposon genes (see above) but also genes with no domain annotation.

After identifying preferred codons, differences in codon usage between different classes of putative effector genes situated on lineage-specific (LS) and core chromosomes were investigated. Two statistics summarising gene codon usage bias were calculated in addition to RSCU and ENc; frequency of optimal codons (Fop) and Codon Bias Index (CBI). Codon usage can be influenced by the selection on the sequence GC content so overall GC content of each gene (GC) and GC content in the third position of synonymous codons (GC3s) were also calculated by codonW. Pairwise correlation between all codon bias metrics, expression level, GC, GC3s content were investigated with Spearman’s rank correlation, and differences in codon usage bias between different subsets of genes compared with a *t*-test. All the statistical analyses were carried out in R.

### Gene Expression

Expression values for predicted genes were determined using Cufflinks to quantify fpkm values of RNAseq reads aligned to the genome during gene prediction. A mean fpkm value was taken for each gene from the three technical replicates. Expression of genes belonging to different effector categories and in different regions of the genome was investigated using negative binomial generalised linear model, with log transformation using the glm function in R and other base functions^57^. A final model tested terms for region (Core, Effector-Rich Core, non-PS LS and PS) gene type (Secreted, CAZyme, EffectorP, non-effector) and whether the gene was within 2 kb of a mimp (Yes, No).

Combinations of terms were combined into a single input factor into the glm. This allowed removal of four combinations that were not present in the dataset, as no CAZyme, EffectorP or Secreted genes were within 2 kb of a mimp and found on core chromosomes, also no secreted genes were within 2 kb of a mimp and were located on effector rich core regions.

## Supplementary Legends

**Supplementary Table 1**

Accession numbers of *Fusarium oxysporum* (Fo) and *F. oxysporum* f. sp *cepae* (Foc) assemblies and gene models deposited at Genbank as Whole Genome Shotgun projects.

**Supplementary Table 2**

Identification of core (C), lineage specific (LS), and pathogen specific (PS) regions in *Fusarium oxysporum* f. sp. *lycopersici* (Fol) genome, through alignment of *F. oxysporum* (Fo) and *F. oxysporum* f. sp. *cepae* (Foc) assemblies. The percentage of non-masked bp covered by aligned sequence is shown for each reference contig.

**Supplementary Table 3**

Distribution of features throughout the *Fusarium oxysporum* f. sp. *cepae* Fus2 genome. Numbers and density of genes predicted as secreted proteins, secreted carbohydrate active enzymes (CAZY), and secreted proteins with an effector-like structure (as predicted by EffectorP). Number and density of secondary metabolite clusters (Sec. Met. clusters) and MIMPs are also shown.

**Supplementary Table 4**

Gene ID and ortholog groups of previously identified *Fusarium oxysporum* f. sp. *lycopersici* (Fol) genes Secreted In Xylem (*SIX*) pathogenicity genes following BLAST searches against *F. oxysporum* f. sp. *cepae* (Foc) Fus2 genome. Genes with no identifiable orthologs are marked as singletons. * Evidence for an additional unpredicted *SIX3* gene was found in the Foc A23 assembly, spanning a contig break.

**Supplementary Table 5**

ID, location, orthogroup, effector evidence, expression in planta and functional annotation for all genes predicted in pathogenic *Fusarium oxysporum* f. sp. *cepae* (Foc) isolate Fus2. Orthogroup contents show the number of proteins clustered into a particular orthogroup by isolate including pathogenic Foc isolates (Fus2, A23, 125), non-pathogenic *F. oxysporum* (Fo) isolates from onion (A13, PG, A28, CB3) and reference genomes for non-pathogenic Fo (isolate fo47) and F. oxysporum f. sp. *lycopersici* (isolate 4287).

**Supplementary Table 6**

SNP variants observed in secreted effector candidates located in pathogen-specific regions of *Fusarium oxysporum* f. sp. *cepae* (Foc) isolate Fus2, through comparison to other Foc isolates.

**Supplementary Table 7**

Comparison of distribution of codon bias across different subsets of genes relative to all genes (t-test), calculated over: all, core, all lineage-specific (LS) chromosomes, LS chromosomes identified as pathogen-specific (PS) and LS chromosomes not identified as PS (non-PS).

**Supplementary Table 8**

Enriched Interproscan terms associated with predicted proteins encoded on(A) pathogen specific (PS) and B) non-PS lineage specific (LS) regions in comparison to core regions. Annotation shown are those significant following Benjamini-Hochberg correction. Terms shown in bold are of particular interest as they were not significantly enriched in other LS regions, or nearly significant (P-values < 0.05 but failed multiple test correction) in other LS regions.

**Supplementary Table 9**

Expression values (fpkm) and annotations of the 50 highest expressed core genes of *F. oxysporum* f. sp. *cepae* Fus2 and those lineage specific (LS) and pathogen-specific (PS) genes showing equal or greater expression.

**Supplementary Table 10**

Identification of transcription factors within the genome of *Fusarium oxysporum* f. sp. *cepae* isolate Fus2 and their expression *in planta*. A) Transcription factors identified in pathogen specific (PS) and non-PS lineage specific (LS) regions. B) Homologs to known *F. oxysporum* f. sp. *lycopersici* transcription factors TF1-9 (B).

**Supplementary Figure 1**

Duplication of genes between and within Fus2 chromosomes show different patterns between core, effector-enriched and LS regions. Links are shown between genes on a Foc contig and all other genes in that orthology group. Genes on core chromosomes (1, 2, 4, 5, 7, 8, 9, 10) do not show high levels of gene duplication across their entire length, with genes in shared ortholog groups primarily on telomeric regions and shared with telomeric regions of other chromosomes. Examples are provided for chromosomes 1 (a) and 4 (b). Duplicated genes on LS regions show a bias towards being shared with other LS regions. Effector enriched core chromosomes (11, 12, 13) show a greater numbers of duplications and are more likely to share genes with one another, as observed in chromosome 11 (c). Genes on Foc LS regions share large numbers of duplicated across their entire length, with duplications primarily shared with other LS regions. Example provided for LS contig 10 (d).

**Supplementary Figure 2**

Repeat masked and gene-coding content of chromosomes and LS regions within Fol and Foc chromosomes / LS regions. Percentage of bp repeat masked in Fol (A) and Foc core (green), non-PS LS (orange) and PS (red) regions (B). Gene density is shown for Fol (C) and Foc (D) core, LS and PS regions. Shaded bars indicate the contribution of genes located in repeatmasked regions towards gene density.

## Author contributions

RJH and JC devised the study. AA carried out genome assembly, annotation, synteny analysis and RNA seq analysis was carried out by SO and LB. AT and AJ carried out the RNA seq experiment, MKS carried out segmental and tandem duplication analysis, variation analysis and codon usage bias studies, BPJG extracted DNA for PacBio sequencing, HB and FW carried out Illumina library preparation and sequencing. AA, AT, MKS, HJB, JC and RJH wrote the manuscript.

## Acknowledgements

The authors wish to thank Hazera Seeds, specifically, Reinout de Heer, Wessel van Leeuwen, Ningwen Zhang, Tosca Ferber and Hans van den Biggelaar for support and advice and David Cole and Cathryn Lambourne for useful discussion. We are grateful to BBSRC (BB/K020870/1 and BB/K020730/1) and AHDB (CP 116) for funding. This article is dedicated to Dr Dez Barbara who was instrumental in the early development of this research.

## Materials and correspondence

For Fusarium isolates and RNA seq data contact Dr John Clarkson. For further information about bioinformatics and sequencing, contact Dr Richard Harrison.

